# Deciphering ocular microbiomes across trachomatous disease severity and *Chlamydia trachomatis* infection using metagenomics

**DOI:** 10.64898/2026.07.16.738988

**Authors:** Augusta Uwamanzu-Nna, Olusola Olagoke, Xiaoyi Charlotte Shi, Hiwot Degineh Mengistie, Kaleb Asfaha, Timothy D. Read, Deborah Dean

## Abstract

*Chlamydia trachomatis* (*Ct*) is the etiologic agent of trachoma, the leading infectious cause of blindness worldwide; ∼100M people are at risk of blindness, particularly in Amhara, Ethiopia, where *Ct* infection and trachoma remain highly prevalent. Currently, there are no data on the role of ocular microbiomes in *Ct* susceptibility and disease pathogenesis. We used metagenomics and multinomial mixture modeling to identify microbial compositions associated with disease severity and *Ct* infection in the microbiomes of 680 Amhara residents. Ten distinct microbial compositions were identified. Healthy microbiomes were dominated by *Mesomycoplasma hyorhinis,* a zoonotic porcine bacterium, and *Staphylococcus aureus*. Both species form biofilms that may competitively prevent or down-regulate chlamydial infections. *Ct*-infected microbiomes had higher species richness and were dominated by *Corynebacterium macginleyi*, *Blautia wexlerae*, a gut microbe associated with dry eye disease, and *S. epidermidis.* The latter two produce acetate that may collectively enhance inflammation and damage mucosal barriers, predisposing individuals to *Ct* infection/persistence. Microbiomes with active inflammation (TF/TI) were dominated by five *Haemophilus* spp., while those with chronic scarring (TS/TT) had low diversity and *Corynebacterium* spp. dominance. *Moraxella* spp. dominated microbiomes with inflammation on-top-of scarring (I-TS/TT) and TS/TT among adult females. Applying our microbiome-based framework to other trachoma-endemic populations will generate further insights into pathogenic and healthy ocular microbiomes on a global scale. Our data highlight discrete microbiome shifts that can be quantified and linked to healthy conjunctivae, *Ct* infection and trachoma severity, forming a foundation for mechanistic and prospective studies to develop prediction models for infection and disease progression.

**IMPORTANCE:** Trachoma, caused by *Chlamydia trachomatis,* is the leading infectious cause of blindness worldwide with over 100M people at risk. Despite decades of the World Health Organization’s SAFE (<u>S</u>urgery, <u>A</u>ntibiotics, <u>F</u>acial cleanliness, <u>E</u>nvironmental improvement) strategy, trachoma remains a public health threat with the highest disease burden in sub-Saharan Africa. Ethiopia is hyperendemic for trachoma even after 15 years of SAFE, indicating an urgent need to study the microbial ecology of ocular microbiomes that may drive infection and disease. Our research is significant in revealing distinct beneficial and pathogenic microbial compositions associated with healthy conjunctivae, *Ct* infection, and mild-to-severe disease, respectively. Beneficial microbes provide the possibility of designing ecology-informed therapeutics as antibiotic alternatives to improve antibiotic stewardship. Moreover, comparing microbial compositions across other trachoma-endemic populations will generate additional global insights into pathogenic and healthy microbiomes that will further inform mechanistic studies and targeted clinical screening to advance the elimination of blinding trachoma.

## INTRODUCTION

Trachoma is the world’s leading infectious cause of blindness with nearly 2M people blind and over 100M at risk(1, 2). Recurrent conjunctival infection with the obligate intracellular bacterium *Chlamydia trachomatis* (*Ct*) from childhood onward can cause chronic disease(3). The socioeconomic consequences of trachoma include billions in lost productivity and reduced quality of life for individuals and their families(1).

The World Health Organization (WHO) developed the SAFE strategy to eradicate blinding trachoma by 2020, a target now extended to 2030(1). SAFE entails <u>S</u>urgery to correct trachomatous trichiasis (TT, in-turned eyelashes), <u>A</u>ntibiotics to treat *Ct* infection, and <u>F</u>acial cleanliness and <u>E</u>nvironmental improvement to prevent infection and reduce *Ct* transmission. Trachoma remains an ongoing public health concern(4). Ethiopia accounts for about 55% of the known global trachoma population(5) with the highest burden in Amhara despite 15 years of SAFE interventions(6, 7). *Ct* infection and disease prevalence remain high, warranting further research into potential drivers of pathogenesis, including chlamydial and other zoonotic microbes, and non-*Ct* commensals and pathogens that promote inflammation and disease progression.

While metagenomic shotgun sequencing (MSS) has been used to study dry-eye disease (DED), blepharitis, keratitis, and meibomian gland dysfunction in developed countries(8), there are no MSS data on microbial compositions of the ocular microbiome to provide insight into *Ct* susceptibility and trachoma pathogenesis. Only culture or 16S rRNA sequencing has been used to describe conjunctival microbiota. One culture study found that *Streptococcus pneumoniae* and *Haemophilus influenzae* were more frequently associated with Ethiopian women than men, and *Ct* infection was more frequent in individuals with TT than those without scarring (TS)(9). In Gambia, children with *S. pneumoniae* and *H. influenzae* were more likely to have trachomatous inflammation-follicular (TF)(10). Studies using 16S rRNA found a higher relative abundance of *Corynebacterium* and *Streptococcus* genera in Gambian adults with TS(11, 12).

Here, metagenomics were used to characterize the microbial diversity and composition associated with healthy conjunctivae, *Ct* infection, and trachomatous disease severity, including inflammation and scarring. Multinomial mixture modeling was used to identify unique microbial clusters among disease and infection states as a microbiome-based framework. We also developed a multi-step workflow to remove likely contaminants and assess retained species’ taxonomic assignments, since this has not previously been performed on low biomass samples like those from the conjunctiva, generating a quality-controlled dataset for downstream analyses.

## MATERIALS AND METHODS

### Study population, trachoma grading, and sampling

The cross-sectional parent study used a non-probability, consecutive sampling method conducted over one year following the last azithromycin mass drug administration (MDA), enrolling 1059 villagers in three regions of Amhara aged 1-99 years(12). The parent study provided trachoma grading confirmed by high-resolution photographs(12,13), metagenomes, and demographic data: age (children, <10 years; adolescents, 10–19 years; adults, 20–49 years; older adults, ≥50 years), sex, employment status, occupation, daily household duties, and village. The WHO simplified trachoma grading(14) included TF, trachomatous inflammation-intense (TI), TS, TT, and concurrent inflammation (TF and/or TI) on-top-of scarring (I-TS/TT). T0 represented absence of clinical signs of trachoma.

### Sample processing, metagenomic shotgun sequencing, and taxonomic profiling

**Figure 1** outlines the methodology pipeline. Per the parent study, conjunctival swabs were treated with a lysozyme cocktail before DNA extraction. Libraries were prepared using Illumina Nextera XT kits and sequenced as 150-bp paired-end reads on an Illumina NovaSeq platform. Raw reads were preprocessed using fastp (v0.23.4)(12,15–17) with human reads depleted using Scrubby(18).

**Figure 1.**
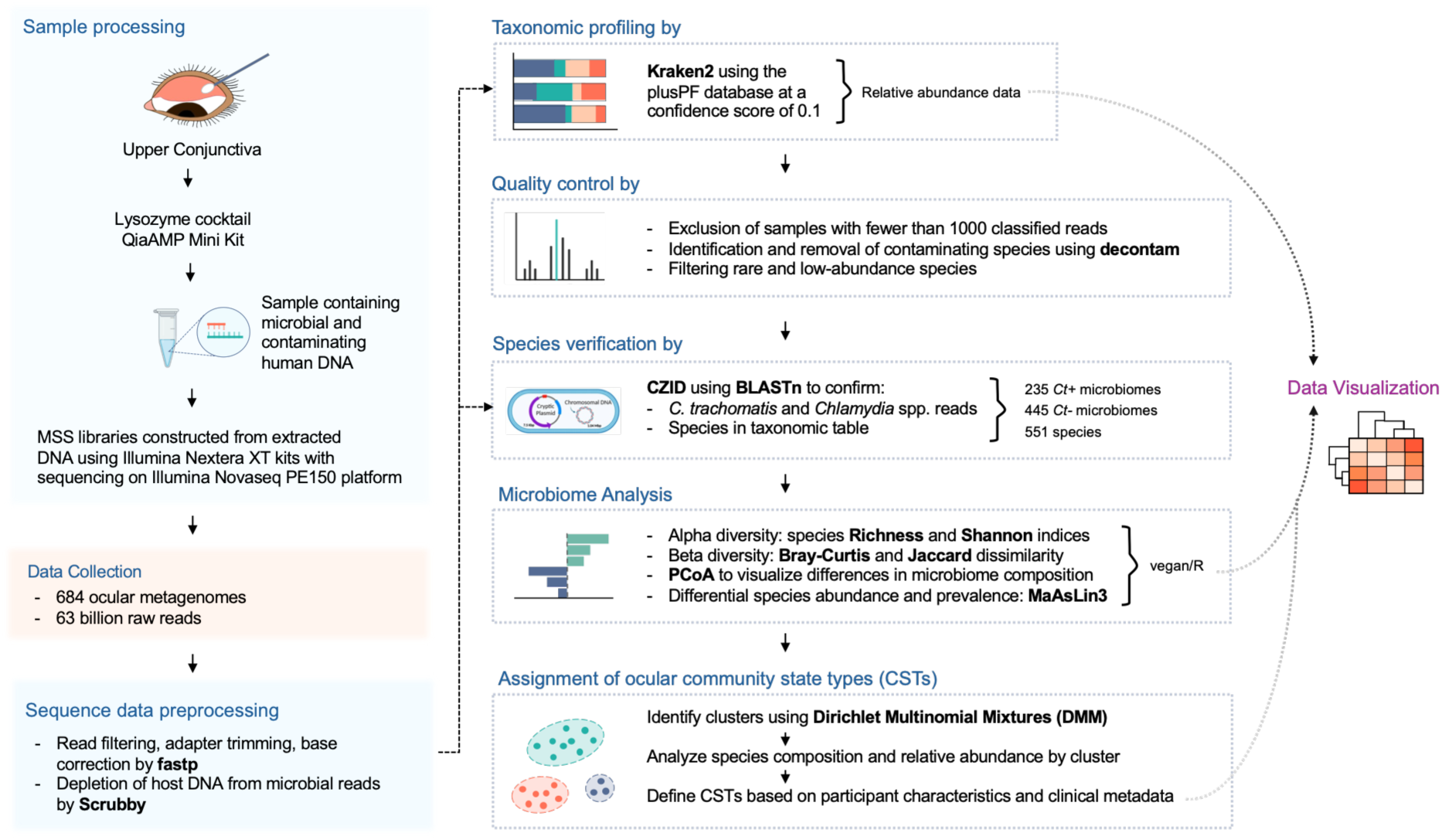
Pipeline for ocular microbiome data generation, preprocessing, and analysis. Adapted from Bommana *et al*.(17).

Kraken2 taxonomic profiling used the plusPF database—combines RefSeq protozoa and fungi genomes—with the Kraken2 Standard database (RefSeq archaea, bacteria, viruses, plasmid, human, and UniVec_Core complete genomes)(19). Kraken2 uses exact k-mer (k=35) matches to classify reads but may generate spurious matches in low-biomass samples like the conjunctiva(20). Low-complexity and homologous sequences can cause multi-mapping, where reads do not align uniquely to one species, and cross-mapping, where reads align to off-target genomes(21). Further, databases that over- or under-represent certain taxa, contain human-contaminated or incomplete genomes, or have inaccurate or sparse taxonomic labels can exacerbate these issues(19,22).

To mitigate these concerns, we applied a Kraken2 confidence score threshold, specifying the minimum proportion of a read’s k-mers mapping to a taxon for assignment at that rank. While higher thresholds can reduce false species identification, unclassified reads can increase(23). To select a threshold, each read classified at the species level under the default confidence setting of 0 received a label score, defined as the number of species-mapping k-mers divided by the total number of unambiguous k-mers in that read(24). Scores were averaged for all reads assigned to each species across all samples, producing a distribution of confidence scores (**Fig. S1).** A score of 0.1 improved read classification precision, and samples with at least 1,000 classified reads were retained.

### Contaminant identification and quality control

Microbial signals in microbiomes may be obscured by DNA contamination from kits, reagents, and/or the environment(25). Decontam (v1.26.0) in R was used to identify and remove contaminating species using the ‘isContaminant’ function, combining frequency and prevalence modes. Frequency (threshold=0.1) identifies species inversely correlated with microbial DNA concentration, while prevalence (threshold=0.2) detects more abundant species in negative controls compared to samples(26). Here, negative controls included nuclease-free water, M4 buffer, control swab elutions, and control swabs in M4 processed alongside samples to detect contaminants during collection, handling, DNA extraction, and/or sequencing. Combining frequency and prevalence methods is optimal for identifying contaminants in low-biomass samples, but prevalence may misclassify genuine taxa shared by clinical samples and controls(20). Additionally, taxa commonly reported as laboratory/reagent contaminants that are environmental may actually comprise conjunctival microbiomes of participants in agrarian settings(27).

To distinguish true microbes from contaminants, we statistically compared species read distributions between samples and controls. Species belonging to genera not previously associated with the eye or unlikely in agrarian regions like Ethiopia were excluded. For species flagged as contaminants but plausibly present in both, we compared the ratio of maximum sample reads to maximum control reads with thresholds based on control read counts. Species ratios exceeding these thresholds were retained, including *Staphylococcus aureus* (<100 maximum control reads; 5x ratio) and *Corynebacterium kefirresidentii* (100-1,000; 10x ratio). Only species with ≥2 reads in ≥10% of samples were retained to limit sequencing artifacts from rare taxa(28). We verified the remaining species using CZ ID (formerly IDSeq) Illumina mNGS Pipeline (v8.3), an open-source platform for pathogen detection in metagenomes(29). Complementing Kraken2’s k-mer approach, it uses minimap2 to directly align reads to reference sequences in a compressed version of NCBI’s NT database (Index Date 2024-02-06). Species coverage was evaluated with built-in tools, and CZ ID integrates with the Basic Local Alignment Search Tool (BLAST) to confirm Kraken2 species identification.

### *C. trachomatis,* zoonotic *Chlamydia* and other non-*Ct Chlamydia* species identification

Samples were screened for *Ct* using in-house quantitative (q)PCR assays targeting single copy *omcB*, *ompA*, and/or multicopy plasmid *pgp*3 genes(16,30). Although highly sensitive and specific for *Ct*, targeting only these genes misses other *Ct* genomic or plasmid regions, potentially underestimating infection prevalence. Although MSS has been used to diagnose infectious diseases(31), low DNA yield limits its effectiveness in low-biomass samples(20). After species identification using Kraken2, microbiomes were queried for *Ct* and zoonotic and non-*Ct Chlamydia* spp. reads using CZ ID and BLAST as described above to confirm species assignment.

*Ct* infection was defined as ≥1 genome copy by qPCR and/or ≥1 MSS read matching the *Ct* genome or plasmid at 99-100%. Similar criteria defined the presence of zoonotic and other non-*Ct Chlamydia* spp.

### Microbial diversity, composition, and differential abundance

To determine population-level changes in conjunctival microbial diversity, observed species counts (i.e., richness) and Shannon Index (i.e., species richness and relative abundance or evenness)(32) were measured for *Ct* and trachoma grade. Differences in species composition were measured using Bray-Curtis and Jaccard beta diversity indices. Bray-Curtis quantifies dissimilarity between samples based on species abundance, while Jaccard is based on species presence or absence(32). Principal Coordinate Analysis (PCoA) was used to visualize group differences.

We used Microbiome Multivariable Associations with Linear Models 3 (MaAsLin 3)(33) to identify associations of species abundance and prevalence with, for example, *Ct* infection while controlling for covariates (e.g., trachoma grade, age group, sex, and sampling depth). MaAsLin 3 accounts for sparsity and compositionality and analyzes absolute and relative abundances. Default settings were used except for a 1% abundance threshold and a 0.05 false discovery rate (FDR) with B-H correction. The same model assessed associations with trachoma grades, sex, and age groups.

### Microbial compositions among ocular microbiomes

Dirichlet Multinomial Mixtures (DMM), a probabilistic model accommodating varying sampling depth, was used to identify differential microbial compositions or Community State Types (CSTs). DMM uses unsupervised learning and an evidence-based framework to determine the optimal number of Dirichlet mixture components, or clusters, that fit a dataset(34). Species-level weight (*π̅*) and precision (*θ̅*) measure size and homogeneity, respectively. CSTs were distinguished by summing absolute differences in mean species relative abundance between each CST and a reference cluster, representing the average microbiome composition across all participants(34). Converted to percentages, sums ranged from 0 (identical) to 200 (completely different). This result quantified each species’ contribution to the calculated differences between CSTs and the reference cluster.

The top 10 species per cluster were assessed using the Dirichlet parameter (*α̅*), a product of that cluster’s precision and taxa relative abundances termed “value.” It reflects how strongly each species contributes to a given cluster. Species composition was evaluated using relative abundance. Shannon indices, age, trachoma grade, *Ct* infection, and the presence of zoonotic and non-*Ct Chlamydia* spp. were compared by CST. We built Firth-penalized logistic regression models for each CST to identify associations between participant characteristics and cluster membership. PCoA was used to visualize CST differences.

### Statistical analysis

Analyses were performed in R using phyloseq (v1.50.0) and vegan (v2.7.1), and plots were generated in R using ggplot2 (v3.5.2) and ggpubr (v0.6.0)(35–38). Alpha diversity comparisons and differences between read distributions for samples and controls were identified using pairwise Wilcoxon Rank-sum tests (B-H correction for multiple comparisons). Shannon Index and age-group differences across CSTs were assessed with the Kruskal-Wallis test, while beta diversity comparisons used Permutational Multivariate Analysis of Variance (PERMANOVA) and Analysis of Similarities (ANOSIM). A *P*<0.05 was considered significant unless otherwise stated.

## RESULTS

### Population characteristics

**Table 1** summarizes bivariate associations between *Ct* infection and age group, sex, trachoma grade, occupation, village region, and zoonotic or non-*Ct Chlamydia* spp. *Ct* prevalence was 22.2% (235/1059). Adults and older adults had significantly lower odds of *Ct* infection than children. Active trachoma was associated with increased odds of *Ct*. Occupations and daily household duties were grouped into seven categories (**Table S1**) with nearly half the population working in agriculture. Pre-school children and students had significantly higher odds of *Ct*. Zoonotic or non-*Ct Chlamydia* spp. occurred in 7.5% (51/680) of the metagenomes (**Table S2**; see *Chlamydia* taxonomy section) and were not significantly associated with *Ct*. Participant metadata and metagenomic data (*N=*684) are in **Table S2**.

**Table 1.**
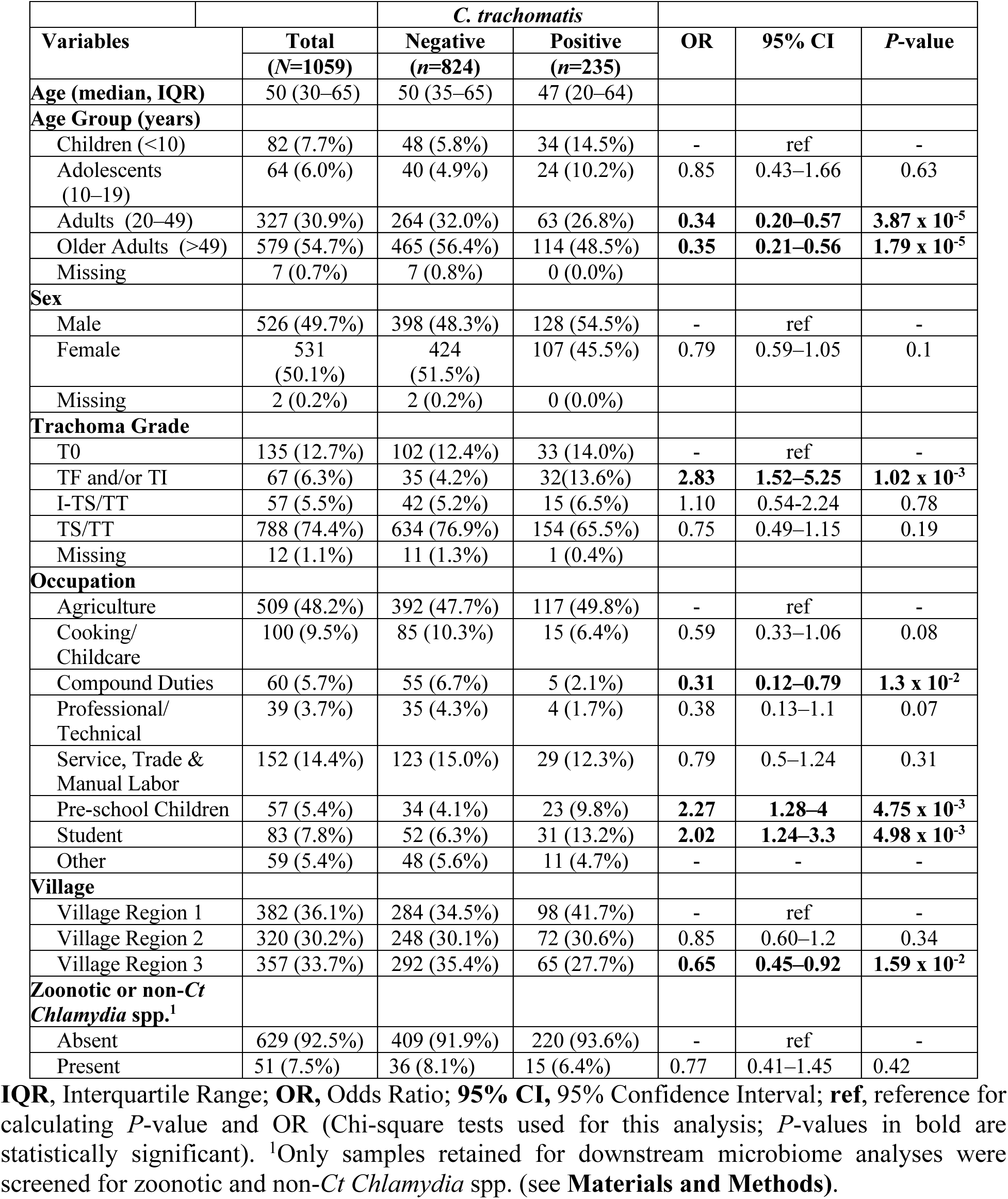
Participant characteristics and bivariate associations of *C. trachomatis* infection with age group, sex, trachoma grade, occupation, and village.

|  |  | *C. trachomatis* |  |  |  |  |
|---|---|---|---|---|---|---|
| **Variables** | **Total** | **Negative** | **Positive** | **OR** | **95% CI** | ***P*-value** |
|  | **(*N*=1059)** | **(*n*=824)** | **(*n*=235)** |  |  |  |
| **Age (median, IQR)** | 50 (30–65) | 50 (35–65) | 47 (20–64) |  |  |  |
| **Age Group (years)** |  |  |  |  |  |  |
| Children (<10) | 82 (7.7%) | 48 (5.8%) | 34 (14.5%) | - | ref | - |
| Adolescents (10–19) | 64 (6.0%) | 40 (4.9%) | 24 (10.2%) | 0.85 | 0.43–1.66 | 0.63 |
| Adults (20–49) | 327 (30.9%) | 264 (32.0%) | 63 (26.8%) | **0.34** | **0.20–0.57** | **3.87 x 10<sup>-5</sup>** |
| Older Adults (>49) | 579 (54.7%) | 465 (56.4%) | 114 (48.5%) | **0.35** | **0.21–0.56** | **1.79 x 10<sup>-5</sup>** |
| Missing | 7 (0.7%) | 7 (0.8%) | 0 (0.0%) |  |  |  |
| **Sex** |  |  |  |  |  |  |
| Male | 526 (49.7%) | 398 (48.3%) | 128 (54.5%) | - | ref | - |
| Female | 531 (50.1%) | 424 (51.5%) | 107 (45.5%) | 0.79 | 0.59–1.05 | 0.1 |
| Missing | 2 (0.2%) | 2 (0.2%) | 0 (0.0%) |  |  |  |
| **Trachoma Grade** |  |  |  |  |  |  |
| T0 | 135 (12.7%) | 102 (12.4%) | 33 (14.0%) | - | ref | - |
| TF and/or TI | 67 (6.3%) | 35 (4.2%) | 32(13.6%) | **2.83** | **1.52–5.25** | **1.02 x 10<sup>-3</sup>** |
| I-TS/TT | 57 (5.5%) | 42 (5.2%) | 15 (6.5%) | 1.10 | 0.54-2.24 | 0.78 |
| TS/TT | 788 (74.4%) | 634 (76.9%) | 154 (65.5%) | 0.75 | 0.49–1.15 | 0.19 |
| Missing | 12 (1.1%) | 11 (1.3%) | 1 (0.4%) |  |  |  |
| **Occupation** |  |  |  |  |  |  |
| Agriculture | 509 (48.2%) | 392 (47.7%) | 117 (49.8%) | - | ref | - |
| Cooking/Childcare | 100 (9.5%) | 85 (10.3%) | 15 (6.4%) | 0.59 | 0.33–1.06 | 0.08 |
| Compound Duties | 60 (5.7%) | 55 (6.7%) | 5 (2.1%) | **0.31** | **0.12–0.79** | **1.3 x 10<sup>-2</sup>** |
| Professional/Technical | 39 (3.7%) | 35 (4.3%) | 4 (1.7%) | 0.38 | 0.13–1.1 | 0.07 |
| Service, Trade & Manual Labor | 152 (14.4%) | 123 (15.0%) | 29 (12.3%) | 0.79 | 0.5–1.24 | 0.31 |
| Pre-school Children | 57 (5.4%) | 34 (4.1%) | 23 (9.8%) | **2.27** | **1.28–4** | **4.75 x 10<sup>-3</sup>** |
| Student | 83 (7.8%) | 52 (6.3%) | 31 (13.2%) | **2.02** | **1.24–3.3** | **4.98 x 10<sup>-3</sup>** |
| Other | 59 (5.4%) | 48 (5.6%) | 11 (4.7%) | - | - | - |
| **Village** |  |  |  |  |  |  |
| Village Region 1 | 382 (36.1%) | 284 (34.5%) | 98 (41.7%) | - | ref | - |
| Village Region 2 | 320 (30.2%) | 248 (30.1%) | 72 (30.6%) | 0.85 | 0.60–1.2 | 0.34 |
| Village Region 3 | 357 (33.7%) | 292 (35.4%) | 65 (27.7%) | **0.65** | **0.45–0.92** | **1.59 x 10<sup>-2</sup>** |
| **Zoonotic or non-*Ct Chlamydia* spp.<sup>1</sup>** |  |  |  |  |  |  |
| Absent | 629 (92.5%) | 409 (91.9%) | 220 (93.6%) | - | ref | - |
| Present | 51 (7.5%) | 36 (8.1%) | 15 (6.4%) | 0.77 | 0.41–1.45 | 0.42 |

**IQR**, Interquartile Range; **OR,** Odds Ratio; **95% CI,** 95% Confidence Interval; **ref**, reference for calculating *P*-value and OR (Chi-square tests used for this analysis; *P*-values in bold are statistically significant). <sup>1</sup>Only samples retained for downstream microbiome analyses were screened for zoonotic and non-*Ct Chlamydia* spp. (see **Materials and Methods)**.

**Table 2** shows a multivariable logistic regression model, adjusted for potential confounders with age group, trachoma grade, occupation, and village as independent variables. Only active trachoma and compound duties remained significant.

**Table 2.** Multivariate associations of *C. trachomatis* infection with age group, trachoma grade, occupation, and village.

| **Variables** | **UOR** | **95% CI** | **AOR** | **95% CI** | ***P*-value** |
|---|---|---|---|---|---|
| **Age Group (years)** |  |  |  |  |  |
| Children (<10) | - | ref | - | ref | - |
| Adolescents (10–19) | 0.85 | 0.43–1.66 | 0.851 | 0.32–2.24 | 0.74 |
| Adults (20–49) | 0.34 | 0.2–0.57 | 0.357 | 0.11–1.21 | 0.10 |
| Older Adults (>49) | 0.35 | 0.21–0.56 | 0.36 | 0.10–1.24 | 0.11 |
| **Trachoma Grade** |  |  |  |  |  |
| T0 | - | ref | - | ref | - |
| TF/TI | **2.83** | **1.52–5.25** | **2.76** | **1.28–5.95** | **9.7 x 10<sup>-3</sup>** |
| I-TS/TT | 1.10 | 0.54-2.24 | 1.72 | 0.76-3.88 | 0.19 |
| TS/TT | 0.75 | 0.49–1.15 | 1.16 | 0.65–2.09 | 0.40 |
| **Occupation** |  |  |  |  |  |
| Agriculture | - | ref | - | ref | - |
| Cooking/Childcare | 0.59 | 0.33–1.06 | 0.554 | 0.31–1.01 | 5.2 x 10<sup>-2</sup> |
| Compound Duties | **0.31** | **0.12–0.79** | **0.303** | **0.12–0.78** | **1.28 x 10<sup>-2</sup>** |
| Professional/Technical | 0.38 | 0.13–1.1 | 0.4 | 0.14–1.17 | 0.09 |
| Service, Trade & Manual Labor | 0.79 | 0.5–1.24 | 0.752 | 0.47–1.20 | 0.23 |
| Pre-school Children | 2.27 | 1.28–4 | 0.486 | 0.12–1.89 | 0.30 |
| Student | 2.02 | 1.24–3.3 | 0.854 | 0.33–2.23 | 0.75 |
| Other | - | - | - | - | - |
| **Village** |  |  |  |  |  |
| Village Region 1 | - | ref | - | ref | - |
| Village Region 2 | 0.85 | 0.60–1.2 | 0.92 | 0.64–1.33 | 0.66 |
| Village Region 3 | 0.65 | 0.45–0.92 | 0.701 | 0.48–1.02 | 0.06 |

**UOR,** Unadjusted Odds Ratio; **AOR,** Adjusted Odds Ratio; **95% CI,** 95% Confidence Interval; **ref**, reference for calculating *P*-value and OR (*P*-values in bold are statistically significant.)

**Figure 2** shows trachoma grade distributions by age and sex. Active trachoma had a bimodal distribution with a dominant peak in young children and a smaller peak in adults; chronic trachoma had a broad distribution, skewed towards older ages. Active trachoma was more prevalent in male than in female children. T0 was concentrated in late adolescence and early adulthood, but with a defined peak among females. I-TS/TT peaked before TS/TT in adult females; in males, both overlapped with a peak later in adulthood. *Ct* infection occurred across all ages with peaks in childhood (both sexes), middle adulthood (females only), and late adulthood (males only).

**Figure 2.**
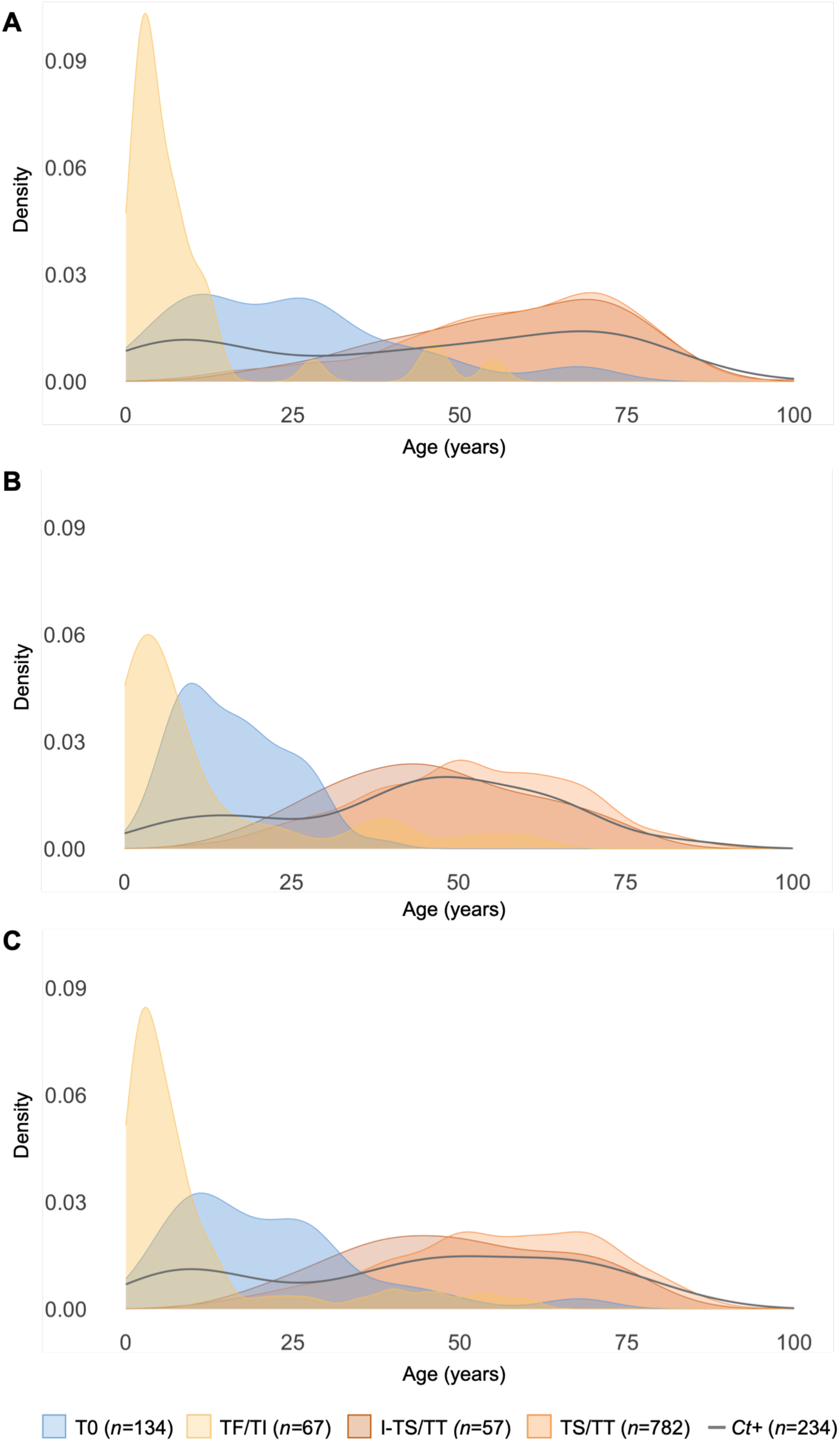
Distribution of trachoma grade among participants by age. **(A)** Males (*n=*516); **(B)** Females (*n=*524); and **(C)** both sexes (*n*=1040). The dark grey line represents the distribution of individuals with *Ct* in the population. Individuals with missing trachoma grade, age, or sex information (*n=*19) were excluded.

### Metagenomic shotgun sequencing results and *Chlamydia* taxonomy

MSS of 684 samples generated ∼63B raw reads, ranging from 6.3M to 348.7M per sample (**Table S3)** with a mean of 98% derived from host DNA. Of 13,687 unique species, 3,235 were flagged as potential contaminants (see **Materials and Methods**) and removed (**Table S4**). Samples with ≥1,000 classified reads were retained (680/684). Removing species without two reads in ≥10% of samples yielded 551 unique species (**Table S4**).

*Ct* assigned reads by CZ ID were identified in 247 of 680 MSSs but did not always align to genomic or plasmid sequences using Megablast. Only 183 (74.1%) of 247 MSSs had confirmed *Ct*-specific genomic or plasmid reads. Of these, 59 (32.2%) were also positive by qPCR. This likely reflects the fact that the qPCR targets only *omp*A, *omc*B and/or plasmid *pgp*3 genes, and no other *Ct* loci in the reads. *Ct* infection was therefore defined as ≥1 genome copy by qPCR and/or ≥1 MSS read with a 99-100% match to the *Ct* genome or plasmid.

Based on trachoma studies in Guinea and Nepal that found zoonotic *Chlamydia* species associated with trachomatous inflammation(13,39,40), we screened each metagenome and identified 51 samples (7.5%) with ≥1 read mapping to these or other non-*Ct Chlamydia* species (**Table S1**), representing species of avian, mammalian or reptilian origin. None were significantly associated with *Ct* infection, trachoma grade, or other participant metadata (**Table S5**).

### Trachomatous scarring is associated with significantly reduced species evenness but not richness, especially among older adults

**Figure 3** shows species alpha diversity by covariates. *Ct* infection was associated with significantly higher species richness with no significant change in the Shannon Index (**Fig. 3A**). Microbiomes with TF/TI did not differ significantly from T0 in Shannon indices or richness (**Fig. 3B**). Both I-TS/TT and TS/TT microbiomes had lower Shannon indices than T0 without significant differences in richness (**Fig. 3B**). TS/TT had significantly higher richness than TF/TI, but the lower Shannon Index was not significant (**Fig. 3B**). Older adults had significantly lower Shannon indices than children, adolescents, and adults with no significant richness changes between groups (**Fig. 3C**). There were no differences by sex (**Fig. 3D**).

**Figure 3.**
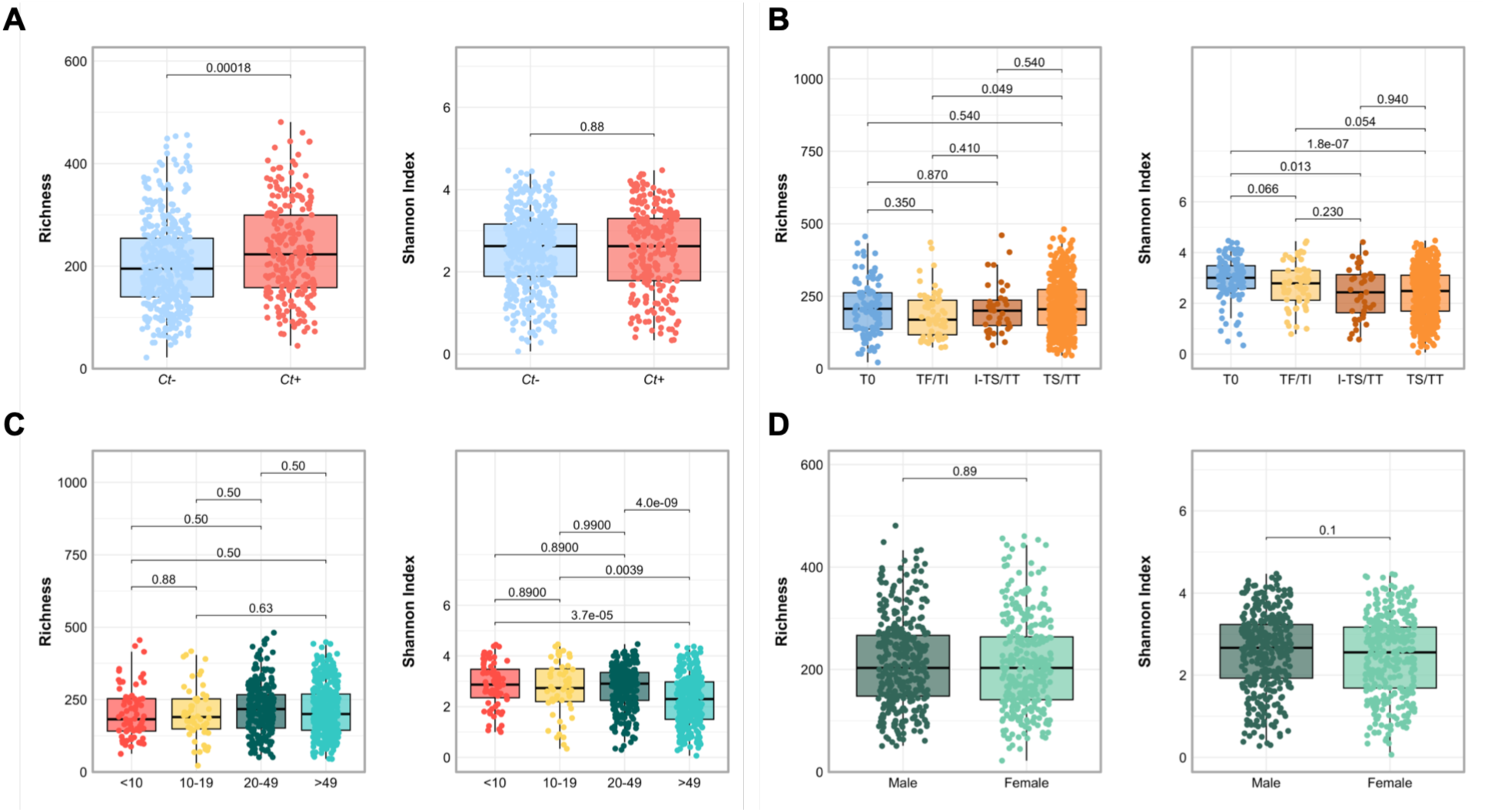
Alpha diversity of ocular microbiomes by *Ct* infection, trachoma grade, age, and sex. Alpha diversity was measured by species richness and the Shannon Index. Box plots display distributions, and *P-*values were computed using the Wilcoxon rank-sum test and adjusted for multiple comparisons using the Benjamini-Hochberg (B-H) method. **(A)** Species richness and Shannon Index for individuals with and without *Ct* infection. **(B)** Species richness and Shannon Index by trachoma grade. **(C)** Species richness and Shannon Index by age group. **(D)** Species richness and Shannon Index by sex.

### Beta diversity is distinct across trachoma grades and age groups

*Ct* and non-*Ct* microbial communities differed significantly by PERMANOVA, but not ANOSIM (**Fig. 4A**), likely reflecting marginal shifts in group centroids despite considerable sample overlap. However, microbiome composition differed significantly by trachoma grade for both tests, although PERMANOVA total variation (*R*^2^) and ANOSIM (*R*) effect sizes were small (**Fig. 4B**). To help explain these differences, Beta diversity analyses were extended to include age group and sex. PERMANOVA detected significant differences by age group and sex for both metrics, while ANOSIM was significant only for age (**Fig. 4C, D**). The similarly low *R*^2^ values across all tests suggest that unmeasured population attributes may explain these differences.

**Figure 4.**
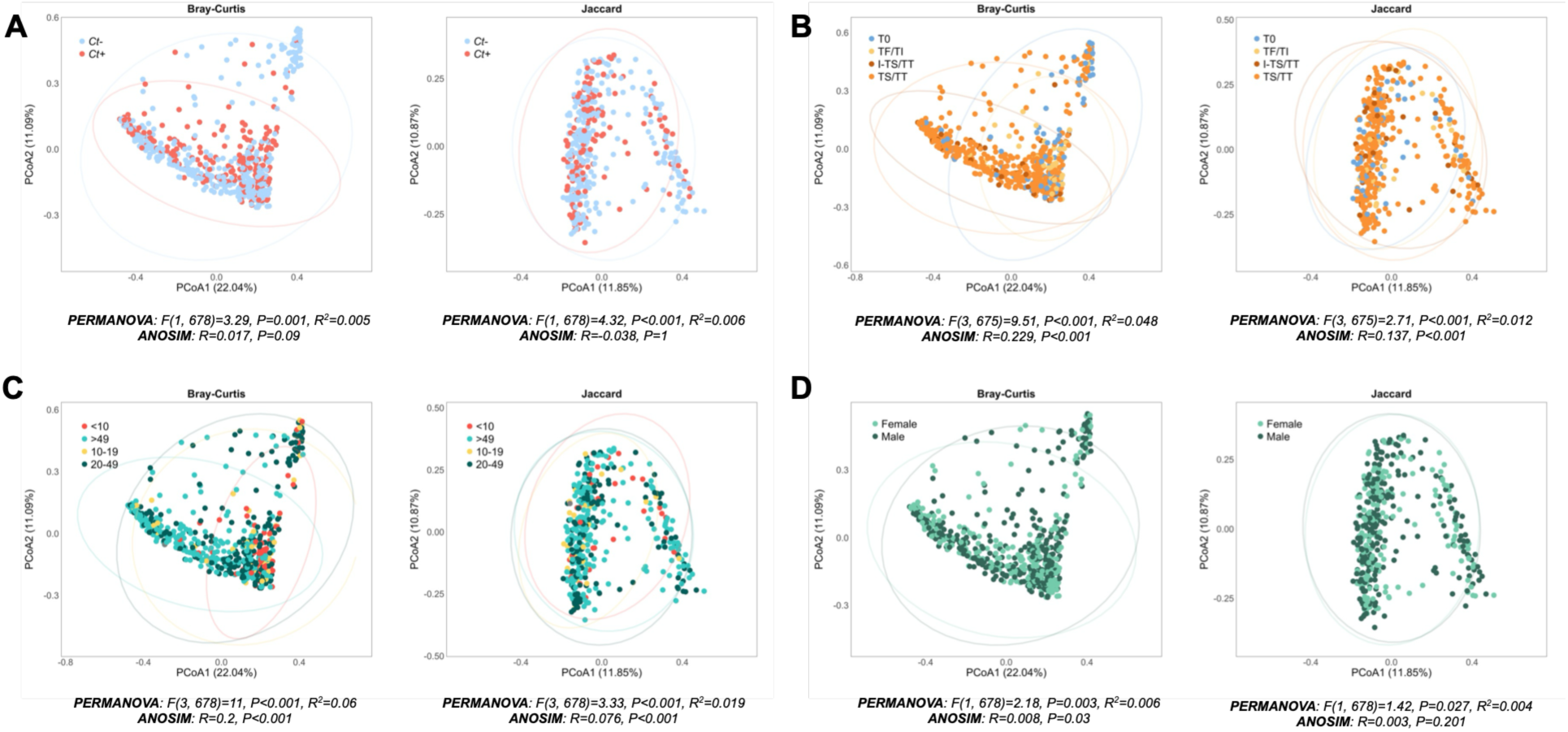
Beta diversity of ocular microbiomes by *Ct* infection, trachoma grade, age, and sex. Beta diversity was assessed using Bray-Curtis and Jaccard dissimilarities and visualized using principal coordinate analysis (PCoA). Group differences were tested using permutational multivariate analysis of variance (PERMANOVA) and analysis of similarities (ANOSIM). Panels show PCoA plots grouped by **(A)** *Ct* infection status (positive or negative), **(B)** trachoma grade (T0, TF/TI, I-TS/TT, and TS/TT), (**C)** age group, and **(D)** sex.

### *Haemophilus* spp. predominate in active trachoma, *Corynebacterium* spp. in chronic trachoma, and *Mesomycoplasma hyorhinis* in healthy *Ct*-negative microbiomes

MaAsLin 3 revealed differentially abundant and prevalent species (**Fig. 5A-E**) with 192 significant associations based on individual or joint *q*-values ≤0.05 (**Table S6**). Most species associated with *Ct* differed in prevalence; only 13 differed in abundance (**Fig. 5A**). *Mesomycoplasma hyorhinis* was the most enriched species in *Ct*-negative microbiomes (**Fig. 5A**). Compared with T0 microbiomes, *Streptomyces subrutilus, H. haemolyticus, H. influenzae,* and *H. parainfluenzae* were significantly more abundant in TF/TI, while *C. macginleyi* and *C. accolens* were most abundant in TS/TT (**Fig. 5B**). I-TS/TT also had a higher *C. macginleyi* and *C. accolens* abundance than T0. Compared with TF/TI, *Moraxella bovis* was the most prevalent and *C. macginleyi* most abundant in I-TS/TT; *S. subrutilus* was the most abundant in TF/TI. Compared with TF/TI, TS/TT had a higher abundance of *C. macginleyi*, *S. epidermidis*, and *C. granulosum*, while *S. subrutilus* and *H. haemolyticus* were enriched in TF/TI (**Fig. 5C**). I-TS/TT and TS/TT microbiomes did not differ significantly.

**Figure 5.**
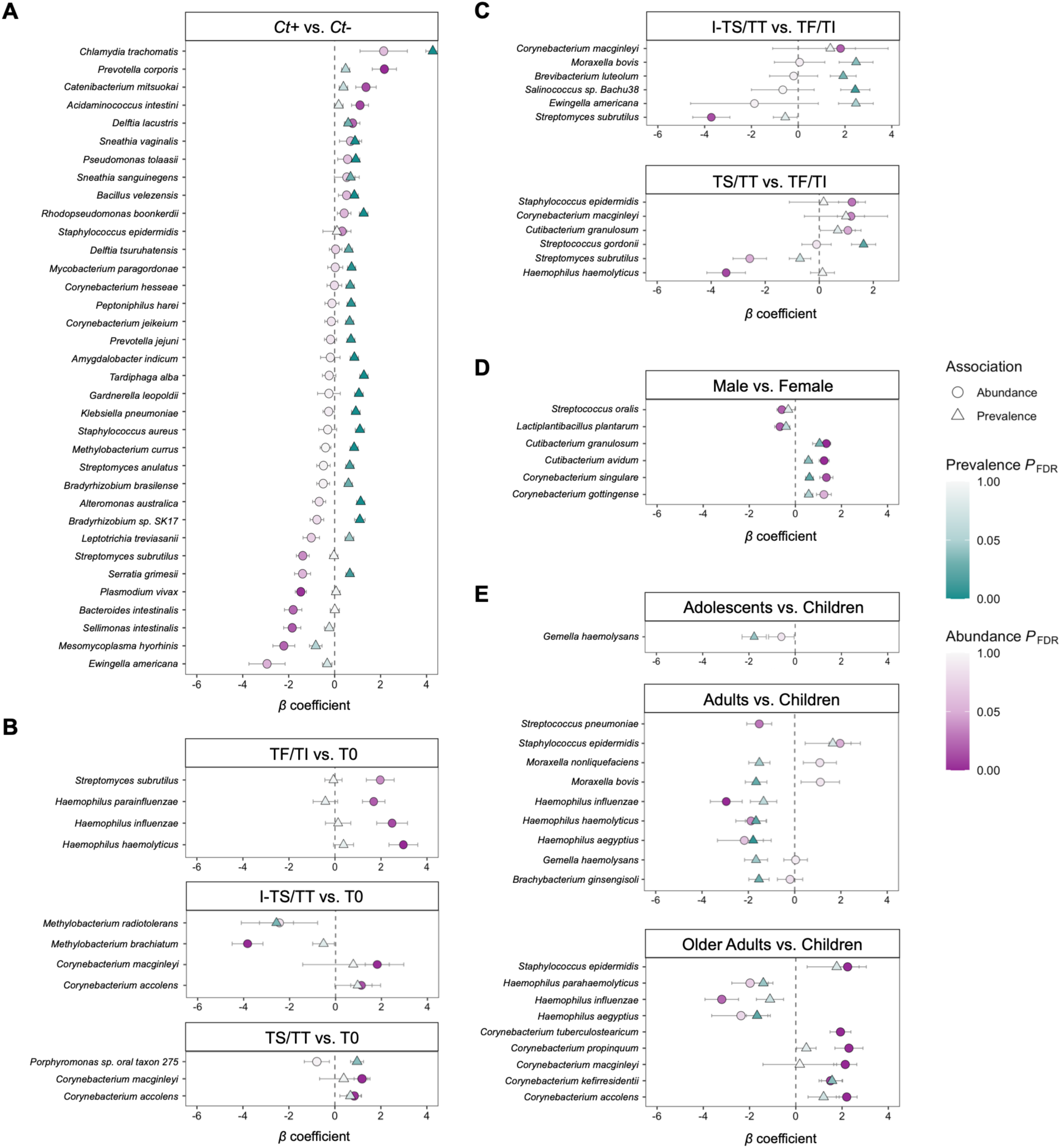
Ocular Microbiome Differential Species Abundance and Prevalence. MaAsLin 3 was used to identify species associated with participant characteristics. Species were ordered based on abundance β coefficients and have either individual or joint (a combination of B-H-corrected abundance and prevalence *P*-values) q-values ≤0.05. For some species, prevalence probabilities could not be computed, so only abundance results are shown. Only significant species are shown, and were identified after controlling for covariates based on **(A)** *Ct* infection (reference: *Ct*-negative); **(B)** active trachoma (TF/TI), inflammation with scarring (I-TS/TT), and chronic trachoma (TS/TT) (reference: T0); **(C)** I-TS/TT, and TS/TT (reference TF/TI); (**D**) sex (reference: females); and **(E)** age group (reference: children).

### Sex and age are associated with distinct microbial profiles

*C. granulosum, C. avidum, C. singular,* and *C. gottingense* were more abundant and prevalent in male than in female microbiomes, whereas *Lactiplantibacillus plantarum* and *S. oralis* dominated female microbiomes (**Fig. 5D**). *H. influenzae* abundance and prevalence were significantly lower in adults and older adults than children and declined with age (**Fig. 5E**; **Fig. S2**). *M. bovis* and *M. nonliquefaciens* were less prevalent but more abundant in adults than children. *S. pneumoniae* was less abundant in adults than children, whereas *S. epidermidis* was more abundant in adults and older adults than children (**Fig. 5E; Fig. S2)**. Five *Corynebacterium* species were more abundant in older adults, but only *C. kefirresidentii* was more prevalent (**Fig. 5E**).

### Distinct microbial compositions are linked to trachoma grades and *Ct* infection

DMM partitioned samples into 10 microbial compositions, largely recapitulating the abundance and compositional patterns above while revealing additional distinctions, providing a microbiome-based framework for characterizing ocular microbial variation in Amhara (**Fig. S3; Fig. 6A**). The top 10 species for each CST are shown in **Figure 6A**. *C. macginleyi* was a top species in six CSTs; CST3 and CST6 had the highest contributions, reflecting species relative abundance and cluster homogeneity (see Materials and **Methods**). *S. epidermidis* also dominated CST3 with the highest mean age and a 95% prevalence of chronic trachoma. CST6 had the highest *C. macginleyi* contribution, five additional *Corynebacterium* species, the lowest mean Shannon Index, and highest chronic trachoma prevalence (96%).

**Figure 6.**
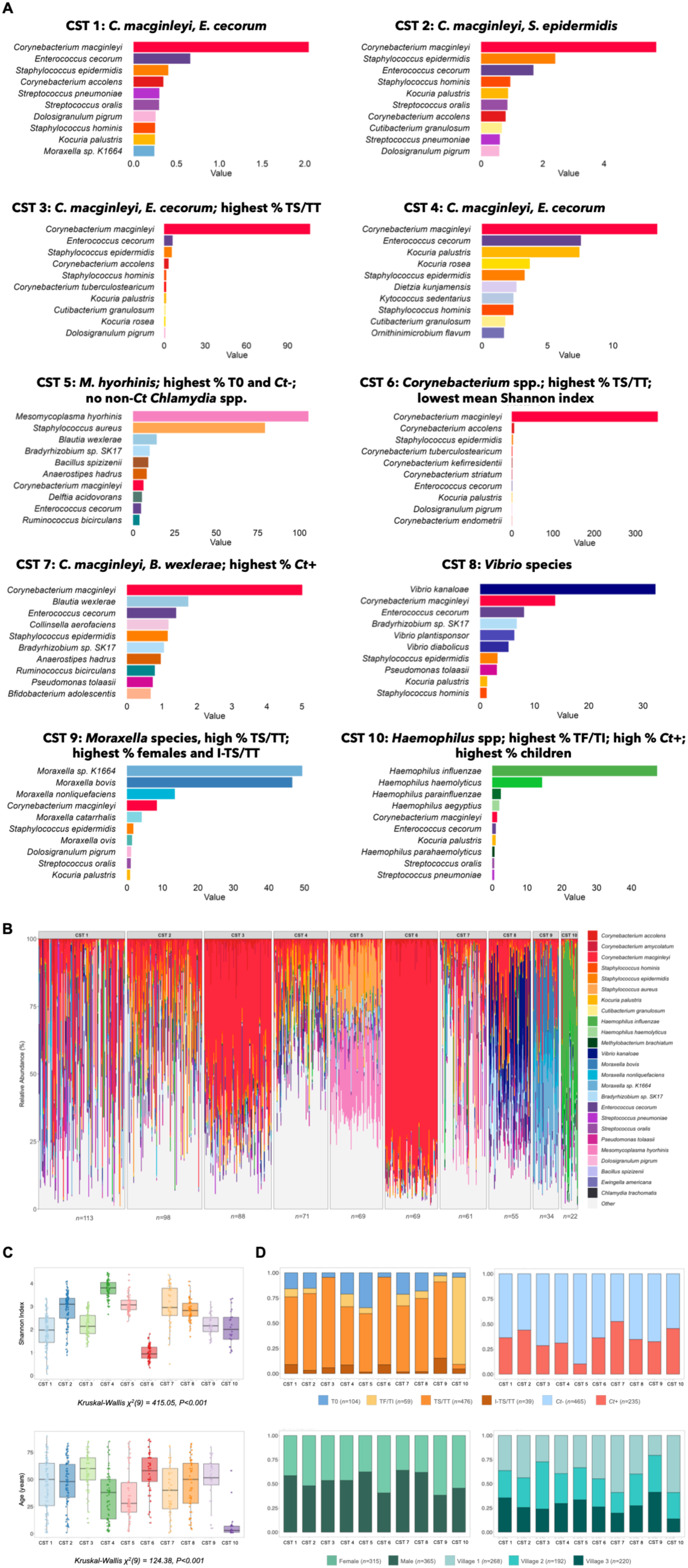
Characteristics of the 10 unique microbial compositions across the ocular microbiomes in Amhara. **(A)** Metagenomes identified microbial compositions; DMM was used to develop CSTs. The top 10 species-level contributions (see **Methods**) for each are shown. Bold headers represent species and villager demographic, clinical, and chlamydial infection characteristics. **(B)** The microbial composition of each sample for each CST using the relative abundance of the top 25 species in that cluster. **(C)** Each CST was analyzed by Shannon Index and age, and the Kruskal-Wallis chi-square test identified significant differences between CSTs**. (D)** The stacked bar plots show the proportions of individuals within each CST stratified by trachoma grade, *Ct* infection status, sex, and village region.

Four clusters had uniquely dominant species. *M. hyorhinis* dominated CST5 with the highest proportion of *Ct-*negative (92%) and T0 (30%) microbiomes, and was the only cluster without zoonotic *Chlamydia* or non-*Ct Chlamydia* spp. (**Table S5**). Three *Vibrio* spp. dominated CST8. Five *Moraxella* spp. dominated CST9 with a high prevalence (91%) of chronic trachoma and the highest proportions of adult and older adult females with I-TS/TT (**Table S7**). CST10 had four dominant *Haemophilus* spp., the highest prevalence of TF/TI (80%) and *Ct* infection (45%), and the lowest mean age. Four remaining clusters had relatively equal species contributions.

**Figure 6B** shows the dominant species distribution within each CST. **Figures 6C-D** show Shannon diversity, age, trachoma grade, *Ct* infection status, sex, and village membership by CST. To evaluate associations between participant metadata and CSTs, we fitted Firth-penalized one-vs-all logistic regression models for each CST, adjusting for covariates (**Table S8**). *Ct*, I-TS/TT, and TS/TT were associated with significantly lower odds of CST5 membership, whereas *Ct* was associated with increased odds of CST7 membership and decreased residence in Villages 2 and 3. Females had much higher odds of CST9 membership, and TF/TI was associated with significantly higher odds of CST10 membership.

### Microbial compositions across the microbiomes highlight distinct profiles in Amhara

Compared with a reference cluster representing the average ocular microbiome of all participants, the most distinct clusters were CST6 (144%), CST5 (132%), CST9 (121%), and CST10 (117%) (**Table S8**). Nine species accounted for 40% of the cumulative differences between CSTs and the reference cluster: *C. macginleyi, H. influenzae, Moraxella sp. K1664, M. bovis, M. hyorhinis, V. kanaloae, S. aureus, H. haemolyticus,* and *E. cecorum* (**Table S10**). CST1 and 2 had the most samples (weight *π̅*_1_ =0.17; *π̅*_2_ =0.15) and highest microbial composition variability (precision *θ̅*_1_ =27.7; *θ̅*_2_ =73.0) (**Table S9**). CST5 and CST6 had the most homogeneous microbiomes (*θ̅*_5_=414.6; *θ̅*_6_=436.2), each capturing 10% (*π̅*_5_=0.10; *π̅*_6_=0.10) (**Table S9**).

**Figure 7** shows the Bray-Curtis PCoA results with 3-D representations highlighting CST clustering (**Fig. 7B-F**; see **Methods**). Microbial composition differed among CSTs with CST5 (salmon), CST6 (red), and CST9 (light purple) showing the best separation.

**Figure 7.**
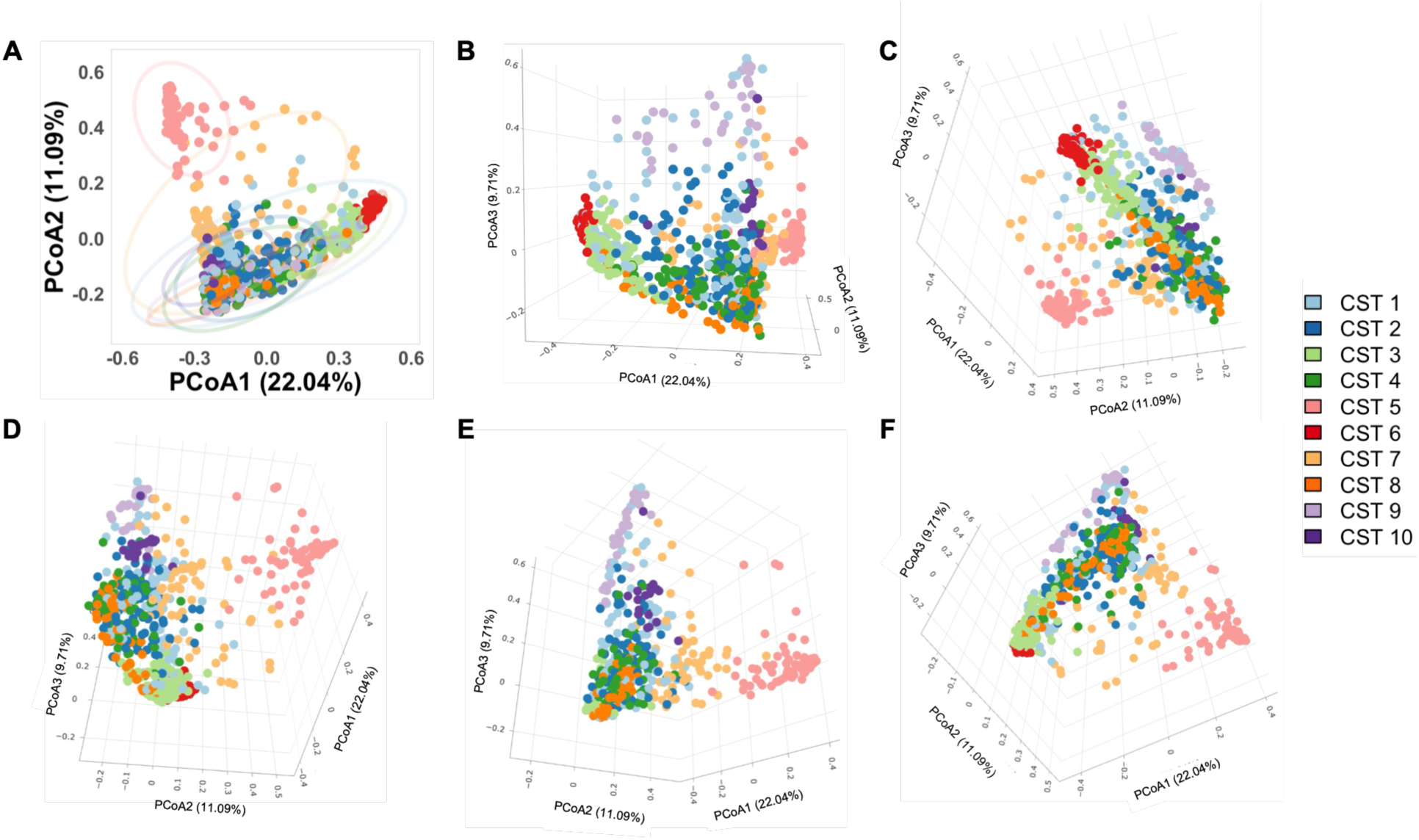
PCoA Representation of CSTs. **(A)** Two-dimensional PCoA plot of Bray-Curtis dissimilarity indices for each CST. Significant differences in microbiome composition were observed using PERMANOVA (F=51.789, *P*<0.001, *R*^2^=0.4103) and ANOSIM (*R*=0.468, *P*<0.001). **(B)** Three-dimensional visualization of CST1 (light blue), CST2 (dark blue) and CST4 (dark green); **(C)** CST3 (light green) and CST6 (red); **(D)** CST5 (salmon) and CST7 (light orange); **(E)** CST9 (light purple) and CST10 (dark purple); and **(F)** CST8 (dark orange).

## DISCUSSION

The global commitment to eliminating blinding trachoma by 2030 requires a fuller understanding of the drivers of *Ct* infection and disease pathogenesis. We evaluated trachoma epidemiology in Amhara, Ethiopia, and used metagenomics and DMM to interrogate microbiomes, providing the largest and most comprehensive assessment of the ocular microbiome in trachoma to date.

Children and those with active trachoma had a higher odds of *Ct* infection, consistent with prior studies identifying them as the primary reservoir of *Ct*(3, 42). Male children had higher *Ct* infection and TF/TI prevalence than female children, reflecting greater organism transfer during close-contact play. Adult females had higher infection, I-TS/TT, and TF/TI prevalence than males and showed scarring nearly two decades earlier (**Fig. 2**). In sub-Saharan Africa, women are the primary caretakers of children, which increases their exposure to *Ct,* potentially explaining their higher I-TS/TT prevalence. This exposure makes them four times more likely than men to develop TS(42,43). Chronic trachoma, resulting from repeated *Ct* infection, inflammation, and progressive scarring(44), was prevalent in older adults. Males had TS/TT at older ages than females, possibly due to increased household contact with infected children once they could no longer work.

These epidemiologic patterns are reflected in ocular microbiome diversity and composition. Although *Ct-*positive microbiomes had higher species richness than *Ct-*negative microbiomes, Shannon diversity did not differ, suggesting the additional taxa were rare and did not alter community evenness. Richness or Shannon indices did not differ between TF/TI and T0, although Shannon diversity tended to be lower in TF/TI (*P*=0.066). A 16S rRNA sequencing study in The Gambia reported reduced richness but unchanged Shannon diversity in TF/TI compared to T0(10). These findings may reflect differences in populations, environmental exposures, and trachoma endemicity, which is low in Gambia. I-TS/TT and TS/TT were associated with significantly lower Shannon indices than in participants without trachoma, especially among older adults, but with unchanged richness, suggesting expansion of a few dominant taxa. Reduced microbial diversity is common in TS(10,11) and a hallmark of ocular surface dysbiosis in DED(45). Children had the highest Shannon diversity, likely reflecting immature hygiene practices, greater physical contact, and exposure to environmental microbes.

The high sensitivity of MSS may amplify contaminants, but there is little consensus on how to address them(46–49). Some studies recommend removing ‘known’ contaminants(25,50,51), but genus-level reporting risks excluding legitimate species(20,52). For example, although *Corynebacterium* spp. have appeared in negative controls(25), several species (e.g., *C. macginleyi, C. accolens, C. striatum*) are frequently isolated from the conjunctiva and implicated in infection and disease(53,54). Our multi-step workflow used decontam to identify contaminants while retaining plausible members of Amhara microbiomes, recognizing that decontam’s prevalence threshold may flag taxa shared by samples and controls(20), some of which may reflect environmental exposures(55). Nearly half of our participants work in agriculture and animal husbandry, and *Bradyrhizobium sp. SK17*, abundant in our dataset, is a common African soil symbiont that fixes atmospheric nitrogen and nodulates legumes(56). Legumes are widely cultivated in Ethiopia for food security, soil fertility, and export(57–59). Although contamination remains possible, agricultural exposure may explain transient or stable presence in ocular microbiomes. Conjunctival *Bradyrhizobium* genera in Gambian villagers have also been attributed to poor housing and sanitation(11). A comparison of hand-associated 16S rRNA sequences found that women from Dar es Salaam, a non-trachoma city, had more soil- and untreated surface water-associated bacteria, including *Bradyrhizobiaceae* and *Rhizobiaceae,* than U.S. women(60). Limited sanitation, safe drinking water, and hygiene(61) likely increase environmental and animal exposures, consistent with our detection of zoonotic and other non-*Ct Chlamydia* spp. of avian, mammalian, and reptilian origin among Amhara residents. These findings highlight the need to expand low-biomass microbiome studies to capture geographic variation driven by environmental and societal factors.

The 10 distinct microbial compositions provide a model-based framework for understanding the microbiome’s role in *Ct* infection and trachoma severity. Overall, healthy microbiomes were dominated by *M. hyorhinis* and *S. aureus*, while more species-rich, *Ct-*infected microbiomes were dominated by *C. macginleyi* and *B. wexlerae*. Diversity and MaAsLin 3 analyses supported a shift from high Shannon diversity and *Haemophilus* spp. dominance in TF/TI to low diversity and *Corynebacterium* spp. dominance in TS/TT. Five *Moraxella* spp. dominated microbiomes in adult and older adult females with I-TS/TT or TS/TT.

A recent 16S rRNA study of healthy European conjunctivae identified nine clusters dominated by *Staphylococcus, Bacillus,* and *Corynebacterium* genera(62). Although *Corynebacterium* may synergize with other commensals to protect the eye from virulent microbes(63), these species also cause conjunctivitis, keratitis, blepharitis, and endophthalmitis(64,65). In our study, *C. macginleyi* was markedly abundant in two clusters (i.e., CST3 and 6), both of which had the oldest participants and highest prevalence of TS/TT (≥95%). TS causes conjunctival goblet cell loss, reduced tear production, and meibomian gland atrophy, promoting DED(66,67). DED is associated with aging, microbiome shifts, and tear film instability from lacrimal or meibomian gland dysfunction (MGD)(68). MGD disrupts the tear film’s outermost lipid layer, potentially favoring lipophilic commensal overgrowth(69). In a 16S rDNA study, *C. accolens-macginleyi* or *Corynebacterium kroppenstedtii* accounted for >75% of reads in 12% of DED patients(65). MGD studies similarly found *C. macginleyi* dominance, highlighting its potential as a disease severity biomarker(70). Three (i.e., *C. accolens, C. tuberculostearicum, C. kefirresidentii*) of five additional *Corynebacterium* species in CST6 are lipophilic like *C. macginleyi*. *C. accolens* is associated with the nares but also DED(64). The latter two produce lipases that suppress other bacteria and facilitate their own growth(65). Expansion of lipophilic microbes like *C. macginleyi* can exacerbate inflammation via increased lipid metabolism(67), supporting their potential role in trachoma pathogenesis.

Common ocular commensals (e.g., *Kocuria palustris, Kocuria rosea*, *Dolosigranulum pigrum, S. epidermidis*) in the clusters are also opportunistic pathogens(71,72). *S. epidermidis* ranked among the top ten species in eight clusters and was differentially abundant in adults, older adults, and participants with chronic trachoma. It predominates in DED(73), where its cholesterol and fatty wax esterases help degrade tear-film lipids(74) and its biofilms promote DED via quorum sensing, virulence factor expression, and inflammation(68), mechanisms that along with *C. macginleyi* could worsen trachoma. Conversely, *S. epidermidis* has probiotic potential—its biofilms and antimicrobial peptides can inhibit opportunists like *S. aureus* through competitive exclusion(75). However, *S. aureus* colonization can dominate niches and displace *S. epidermidis* and other protective commensals, as shown in the skin(76). Consistent with this, *S. aureus* occurred at a high relative abundance without *S. epidermidis* but only in the ‘healthy’ CST5, which had the highest proportion of *Ct-*negative and T0 individuals and uniquely lacked zoonotic and non-*Ct Chlamydia* spp. (**Fig. 6**), suggesting interspecies competition or inhibition.

*M. hyorhinis,* a porcine respiratory commensal capable of systemic infection in pigs(77), was the most abundant bacterium in the healthy cluster. Pig production, an emerging export sector in Amhara, typically involves adults aged 21-39 years(78). In China, which produces and consumes roughly half the world’s pigs, *M. hyorhinis* was enriched in the ocular microbiomes of young adults versus older adults (mean: 27.9 versus 67.1 years)(79,80), suggesting porcine husbandry as a zoonotic source. Like *S. aureus, M. hyorhinis* may initiate biofilms(81,82), thereby preventing *Ct* and/or other *Chlamydia* spp. infections. *Mycoplasma* spp., including *M. hyorhinis*, can inhibit *Ct* and *C. pneumoniae* infectivity and growth *in vitro*(*83*), although urogenital *Ct* strains frequently co-exist with *M. genitalium* in the vagina(84). These findings warrant mechanistic studies to further explore interactions among *M. hyorhinis*, *S. aureus, S. epidermidis, Ct*, and other *Chlamydia* spp.

Three *Vibrio* spp., including the opportunistic aquacultural pathogens *V. kanaloae* and *V. diabolicus*(*85*), dominated one cluster. *Vibrio* spp. have been isolated from Lake Tana, a major Amhara fishing site, despite contamination by fertilizers, municipal sewage, and pit latrines(86); *V. plantisponsor* is associated with wild rice(87). Other zoonotic and environmental microbes among the clusters included *E. cecorum*, an emerging poultry pathogen(88); *Ornithinimicrobium flavum,* a plant microbe(89); and marine *Kytococcus sedentarius,* an opportunistic human pathogen(90).

*H. influenzae,* a leading cause of childhood conjunctivitis, and three other *Haemophilus* spp. were most abundant among children, those with TF/TI, and *Ct-*infected participants in CST10. *H. influenzae* and *Streptococcus* spp. frequently occur in active trachoma(9), and *S. pneumoniae* was significantly more abundant in children. These nasopharyngeal colonizers can reach the eye via fomites or autoinoculation.

*C. macginleyi* and *B. wexlerae,* a gut microbe associated with DED(73), dominated *Ct-*positive microbiomes clustering in CST7. *B. wexlerae* produces acetate, which suppresses NF-κB activity and inflammation at low levels but can be proinflammatory in excess(91). Excess acetate has also been documented in *S. epidermidis*, another CST7 member, and in *C. acnes* infections of the skin(91). Because acetate requires no receptor uptake(91), it spreads regionally, potentially increasing ocular concentrations from adjacent skin or nasopharyngeal colonization. Together, *B. wexlerae* and *S. epidermidis* may promote inflammation and disrupt mucosal barriers, predisposing to *Ct* infection, persistence, and tissue spread. These host-pathogen interactions require further study.

Five *Moraxella* spp. were dominant in microbiomes mostly among adult and older adult females who had the highest proportion of I-TS/TT, clustering in CST9. These species elicit proinflammatory responses that contribute to trachomatous disease(92,93). Prior studies found *Moraxella* prevalent in Ethiopian individuals with TT(8,94). *M. catarrhalis* commonly colonizes children’s conjunctivae and nasopharynx and can cause conjunctivitis(95). Concurrent *Moraxella* infection and I-TS/TT, a rarely reported grade(96,97), suggest that repeated exposure during childcare may increase female caregivers’ infection risk, as with *Ct*.

There are some limitations to our work. Our cross-sectional study cannot capture microbial shifts such as those accompanying progression from repeated TI to TS that occur over years or decades. Further, metagenomics of other trachoma-endemic populations could elucidate similar or distinct microbial profiles to refine commensal, pathogenic and beneficial microbe associations with ocular health and disease. Finally, our inferences about acetate-driven inflammation and biofilm-mediated competition rely on species-level taxonomy and the literature and therefore further mechanistic validation should be considered.

Our findings represent a significant contribution to our understanding of the ocular microbiome in trachoma and disease pathogenesis. Metagenomics revealed that microbiomes with *Ct*, inflammation, inflammation with scarring, and scarring alone each have distinct pathogenic compositions, while healthy conjunctivae have beneficial microbes. The latter provides the possibility of developing ecology-informed therapeutics as alternatives to antibiotics to improve antibiotic stewardship. These data highlight discrete microbiome shifts that can be quantified and linked to *Ct* infection and trachoma severity, forming a foundation for future mechanistic and prospective studies to develop prediction models for infection and disease progression. Further, applying our microbiome-based framework to other trachoma-endemic populations will provide insights into pathogenic and protective microbes on a global scale. Our findings and approach along with targeted clinical screening will help advance the global elimination of blinding trachoma.

## ACKNOWLEDGEMENTS

We wish to thank the villagers of Amhara, Ethiopia, and the Bahir Dar Specialty Eye Center healthcare providers for their assistance with field work and dedication to ocular health care.

A.U.: methodology, formal analysis, investigation, software, validation, data visualization, writing—original manuscript, review and editing.

O.O.: data curation, methodology, formal analysis, investigation, software, validation, supervision, writing—review and editing.

X.C.S.: methodology, formal analysis, writing—review and editing. H.W.D.: methodology, supervision, writing—review and editing.

K.A.: methodology, supervision, writing—review and editing.

T.D.R.: conceptualization, methodology, investigation, funding acquisition, resources, and writing—review and editing.

D.D.: conceptualization, methodology, formal analysis, investigation, funding acquisition, resources, project administration, supervision, data visualization, writing—original manuscript, review and editing.

## SUPPLEMENTAL MATERIAL

### Supplementary Figures

**Figure S1. Kraken2 confidence score analysis.** Kraken2 was used to map metagenomic reads to species using default settings. For each species, per-read confidence scores were calculated as the ratio of k-mers assigned to each species to the total unambiguous k-mers in the read. The distribution of scores is shown, and a confidence score threshold of 0.1 was selected.

**Figure S2: MaAsLin 3 linear associations for differentially abundant species by age group.** MaAsLin 3 was used to identify differentially abundant species based on model variables of *Ct* infection, trachoma grade, age group, sex, and the number of classified reads per sample. Linear association plots were generated for significant species, and differences in species abundance by age group are shown.

**Figure S3. Dirichlet Multinomial Mixtures (DMM) model fit.** The optimal number of Dirichlet mixture components (clusters) that fit this study’s dataset was determined using the Laplace approximation to the negative log model evidence.

### Supplementary Tables

**Table S1.** Occupation categories based on reported employment and daily household duties.

**Table S2.** Participant metadata.

**Table S3**. Metagenomic shotgun sequencing results and quality control statistics.

**Table S4**. Taxonomic profiling, contaminant identification, and quality control pipeline.

**Table S5**. Prevalence of non-*Ct Chlamydia* species by CST grouped and species origin (avian, mammalian, or reptilian).

**Table S6**. Significant species associations based on MaAsLin 3 abundance and prevalence methods.

**Table S7**. Data Values for Figures 1, 2, 5C and 5D.

**Table S8**. Firth-penalized logistic regression model of participant metadata and ocular CST membership.

**Table S9.** Characteristics of the clusters from the DMM output.

**Table S10**. Mean species abundances (%) ranked by contributions to differences between mixture and reference components.

## DATA AVAILABILITY

Pre-processed FASTQ files devoid of human reads from the metagenomic data are available at NCBI-SRA under BioProject PRJNA1440961. Data for reconstructing Figures 1, 2, 5C, and 5D are available in **Table S7**.

## ETHICS APPROVAL

The Institutional Review Boards of UCSF and the Ethiopian Ministry of Health approved in accordance with the Declaration of Helsinki. Written informed consent was provided by all participants except that parents gave written consent for minors. Consent was administered by the Medical Staff from Bahir Dar Eye Specialty Center who have knowledge of the local language and customs. All clinical and demographic data were provided de-identified with no trace to patient names by the parent study to this study for the analyses presented here(12).

