## Supplementary Figures S1-S3 for "Deciphering ocular microbiomes across trachomatous disease severity and *Chlamydia trachomatis* infection using metagenomics"

**Figure S1. Kraken2 confidence score analysis.** Kraken2 was used to map metagenomic reads to species using default settings. For each species, per-read confidence scores were calculated as the ratio of k-mers assigned to each species to the total unambiguous k-mers in the read. The distribution of scores is shown, and a confidence score threshold of 0.1 was selected.

**Figure S2: MaAsLin 3 linear associations for differentially abundant species by age group.** MaAsLin 3 was used to identify differentially abundant species based on model variables of *Ct* infection, trachoma grade, age group, sex, and the number of classified reads per sample. Linear association plots were generated for significant species, and differences in species abundance by age group are shown.

**Figure S3. Dirichlet Multinomial Mixtures (DMM) model fit.** The optimal number of Dirichlet mixture components (clusters) that fit this study's dataset was determined using the Laplace approximation to the negative log model evidence.

### Supplementary Tables

**Table S1.** Occupation categories based on reported employment and daily household duties.
**Table S2.** Participant metadata.
**Table S3**. Metagenomic shotgun sequencing results and quality control statistics.

**Table S4**. Taxonomic profiling, contaminant identification, and quality control pipeline.
**Table S5**. Prevalence of non-*Ct Chlamydia* species by CST grouped and species origin (avian, mammalian, or reptilian).
**Table S6**. Significant species associations based on MaAsLin 3 abundance and prevalence methods.
**Table S7**. Data Values for Figures 1, 2, and 5C and 5D.
**Table S8**. Firth-penalized logistic regression model of participant metadata and ocular CST membership.
**Table S9.** Characteristics of the clusters from the DMM output.
**Table S10**. Mean species abundances (%) ranked by contributions to differences between mixture and reference components.

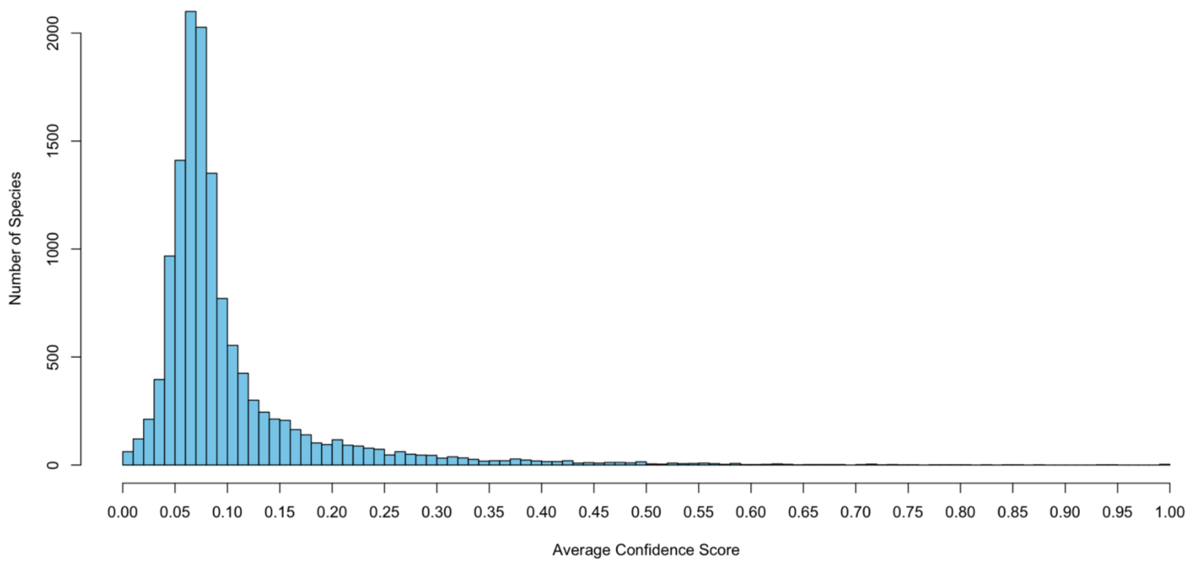


**Figure S1. Kraken2 confidence score analysis.** Kraken2 was used to map metagenomic reads to species using default settings. For each species, per-read confidence scores were calculated as the ratio of k-mers assigned to each species to the total unambiguous k-mers in the read. The distribution of scores is shown, and a confidence score threshold of 0.1 was selected.

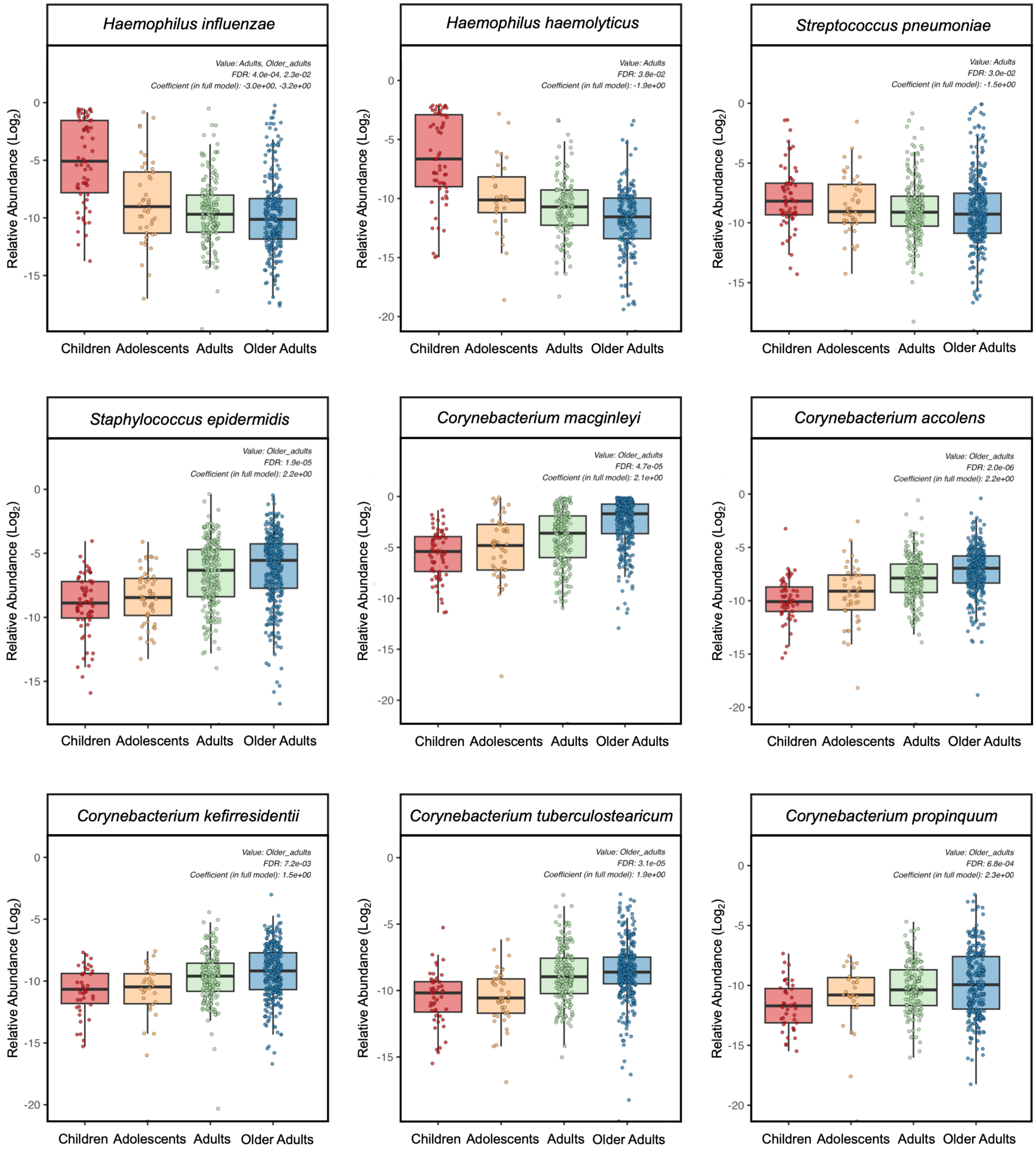


**Figure S2: MaAsLin 3 linear associations for differentially abundant species by age group.** MaAsLin 3 was used to identify differentially abundant species based on model variables of *Ct* infection, trachoma grade, age group, sex and the number of classified reads per sample. Linear association plots were generated for significant species, and differences in species abundance by age group are shown.

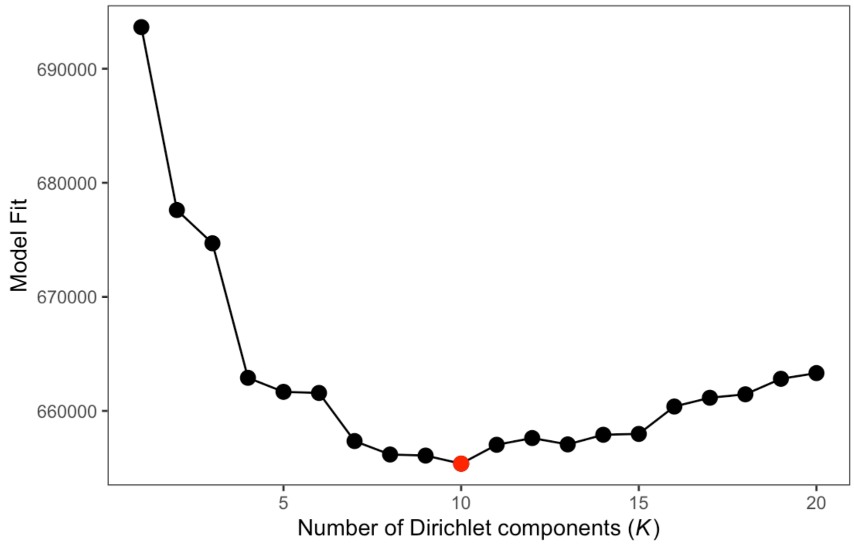


**Figure S3. Dirichlet Multinomial Mixtures (DMM) model fit.** The optimal number of Dirichlet mixture components (clusters) that fit this study's dataset was determined using the Laplace approximation to the negative log model evidence.
